# Phytochemical screening of *Colocasia gigantea* and *Colocasia affinis* (Family: Araceae) using ^1^H-NMR and ^13^C-NMR techniques

**DOI:** 10.1101/2020.10.27.357590

**Authors:** Safaet Alam, Mohammad Rashedul Haque

## Abstract

*Colocasia affinis* and *Colocasia gigantea* are two of commonly found species under *Colocasia* genus. Folkloric use of these plants ascertains their ethnopharmacological importance and these plants are eaten as vegetables in several regions all around the world while *Colocasia gigantea* has planted as an ornamental plant too. Phytochemical screening of dichloromethane fractions of these plants using several separation techniques along followed by ^1^H-NMR and ^13^C-NMR techniques provide flavonoids and some other phyto compounds. However, in total 7 compounds were isolated from these plants i.e. penduletin (1), 7,8 -(3”,3”-dimethyl-pyrano)-4’-hydroxy flavonol (2), mixture of 7,8 -(3’’,3’’-dimethyl-pyrano)-4’-hydroxy flavonol from *C. affinis* and mixture of α-amyrin and β-amyrin (3), penduletin (5), monoglyceride of stearic acid (6) from *C. gigantea*.

## Introduction

Since time immemorial the history of humankind has availed plant extracts of various plants to treat and heal their physical agony (1) and from that ancient time plant and its extracts play a pivotal role and confers therapeutic importance in world health care up to today (2). The gravity of the plant extract resides on the fact that 80% of drug substances are either direct derivation of natural component or a developed version of those natural moieties (3). *Colocasia* leaves have demonstrated the ability of antidiabetic, antihypertensive, immunoprotective, neuroprotective and anticarcinogenic activities. The detailed assessment of phytochemical compounds present in various extracts of the leaves shows the presence of active chemical compounds like anthraquinones, apigenin, catechins, cinnamic acid derivatives, vitexin, and isovitexin which are possibly responsible for the exhibited biological properties (4).

The genus *Colocasia* is represented by 13 species worldwide (5) among which eight species were found in Asia and the Malay Archipelago initially (6). In Bangladesh, so far this genus contains the following nine species: *C. gigantea* (Blume) Hook. f., *C. fallax* Schott, C. *affinis* Schott, *C. esculenta* (L.) Schott, *C. oresbia* A. Hay, *C. heterochroma* H. Li et Z.X. & Wei*, C. virosa* Kunth, *C. lihengiae* C.L. Long et K.M. Liu, and *C. mannii* Hook. f. (7). *Colocasia* is a flowering plant genus under Araceae family native to southeastern Asia and the Indian subcontinent which are widely cultivated and naturalized in other tropical and subtropical regions (8). Genus of *Colocasia* leaves has demonstrated the potentiality of antidiabetic, antihypertensive, immunoprotective, neuroprotective, and anticarcinogenic activities (9). Phytochemical extraction and structure elucidation of *Colocasia* leaves yield chemical compounds such as isoorientin, orientin, isoschaftoside, Lut-6-C-Hex-8-C-Pent, vicenin, alpha-amyrin, beta-amyrin, mono glycerol stearic acid, penduletin anthraquinones, apigenin, catechins, cinnamic acid derivatives, vitexin, and isovitexin (9, 10)

*Colocasia affinis* belongs to Araceae family is a perennial herb of 46 to 91 cm long and can be found all over in Bangladesh (11). In Bangladesh locality, his plant is locally known as Kochu which is also familiar as Dwarf elephant’s ear. In local community the plant leaves are believed effective in cataract treatment and also used as an iron rich soup boiled with coconut milk. Previous study has reported that this plant has antioxidant, cytotoxic, antimicrobial, analgesic, antiinflammatory and dose dependent antidiarrheal activity (11). *Colocasia gigantea* is another species under this genus which generally grows in Thailand and also some southeast Asian countries (12). Tubers and leaf stalk of this plant is consumed in the Pacific islands similar to the people of India and Bangladesh (12–14). Besides, “Bon curry”, a thai food is prepared using stem of *C. gigantea. C. gigantea* is also used as folkloric medicine in Thailand to treat fever, stomach discomfort, infections, drowsiness and to get rid of phlegm. Earlier research has documented anticancer activity of tuber part of *C. gigantea* (15). The present work was carried on focusing the isolation and identification of bioactive secondary metabolites from these two plants.

## Materials and Method

### General experimental procedures

Solvent-solvent partitioning was done by using the protocol designed by Kupchan and modified by Van Wagenen *et al*., 1993 (16) to avail four fractions i.e. n-hexane, dichloromethane, ethyl acetate and aqueous fraction. The ^1^H-NMR and ^13^C-NMR techniques were performed on a Bruker VNMRS 500 instrument using CDCl_3_ as solvent and the chemical shifts were documented in ppm keeping TMS as reference. Gel permeation chromatography (GPC/SEC) was carried over Sephadex (LH-20) (Sigma-Aldrich) along with PTLC and TLC conducted on a silica gel 60 F254 on aluminum sheets at a thickness of 0.25 mm (Merck, Germany) which were observed under UV lamp (UVGL-58, USA) at 254 and 365 nm. Visualization of developed plates were done after spraying vanillin sulfuric acid mixture followed by heating for 5 minutes at 100° C.

### Collection, identification and extraction of plant material

The leaves of *C. affinis* and *C. gigantea* were collected from Moulovibazar and Bandarban, Bangladesh in May 2019. The plants were identified by the experts of Bangladesh National Herbarium, Mirpur, Dhaka and voucher specimens (DACB; Accession no: 57065 and 57066 respectively) have been deposited for these collections. With a proper wash, the whole plants were sun-dried for several days to make it dried followed by grounding these to coarse powders by using a high capacity grinding machine separately. 800 gm of the powdered material was of each plats were taken in two clean, and round bottom flasks (5 liters) and soaked in 2.4 liters of methanol individually. The containers were kept for a period of 30 days along with daily shaking and stirring. The whole mixture acquired from two containers were then filtered through fresh cotton plug and finally with a Whatman No.1 filter paper. The volume of two filtrates were then reduced through using a Buchi Rotavapor at low temperature and pressure. The weight of the crude extract from *C. affinis* and *C. gigantea* were found 71.25 gm and 60.82 gm respectively.

**Figure 1:**
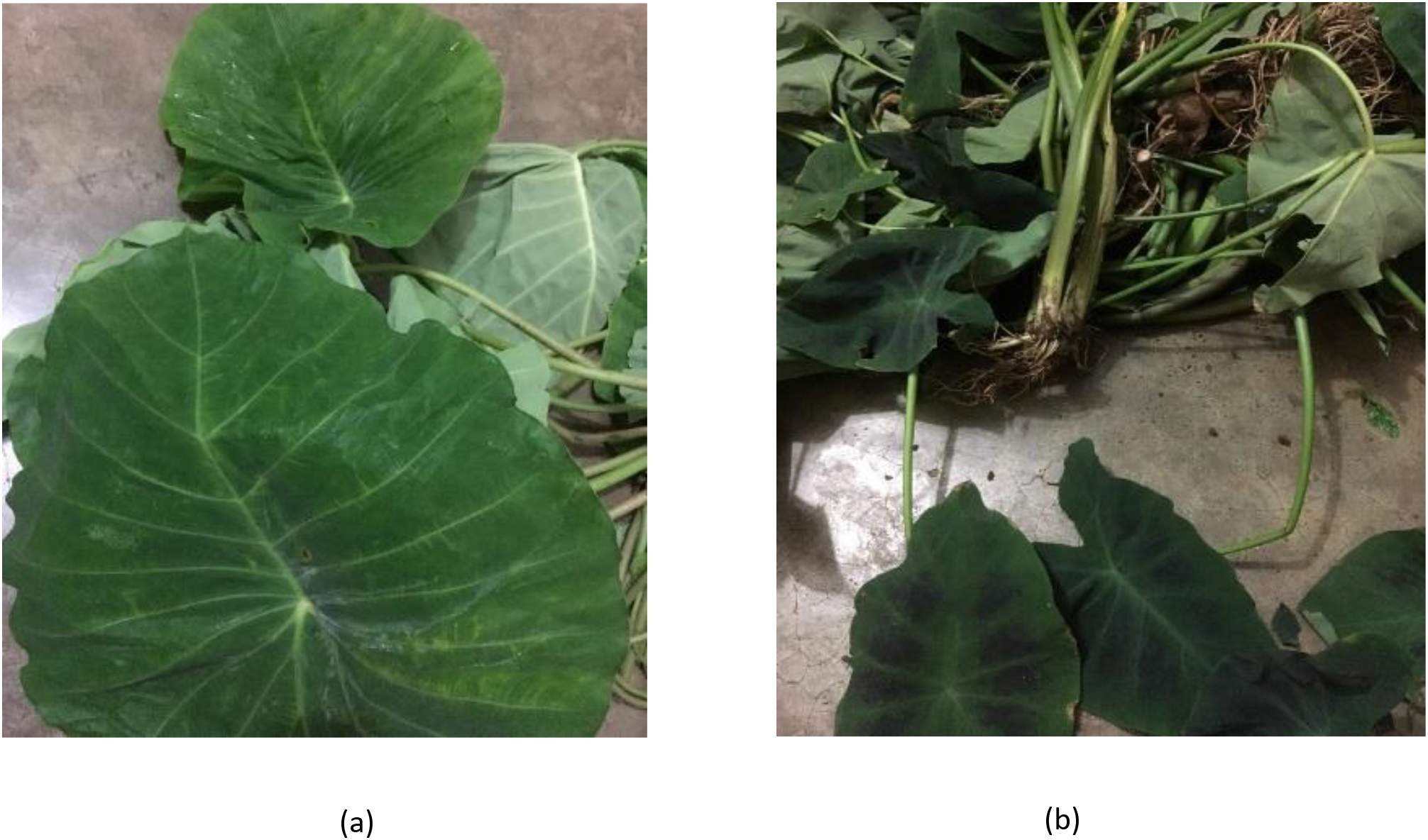
Images of *Colocasia gigantea* (a) and *Colocasia affinis* (b)

### Properties of isolated compounds

#### Penduletin (1 and 5)

Pale yellow crystals, ^1^H NMR (500 MHz, CDCl_3_): δ_h_ 6.59 (s, H-8), 7.90 (d, *J*= 7.90, H-2’), 7.04 (d, *J*=7.2, H-3’), 7.04 (d, 7.2, H-5’), 7.90 (d, *J*=7.2, H-6’), 3.90 (s, 3-Ome), 4.02 (s, 6-Ome), 4.04 (s, 7-Ome), 12.79 (s, 5-OH), 6.37 (s, 4’-OH)

^13^C NMR (500 MHz, CDCl_3_): δ_C_ 148.64, 139.68, 186.19, 159.34, 157.19, 103.75, 123.50, 128.04, 114.61, 162.68, 114.61, 128.04, 61.85, 55.56

#### 7,8 -(3”,3”-dimethyl-pyrano)-4’-hydroxy flavonol (2)

White powder, ^1^H NMR (500 MHz, CDCl_3_): δ_H_ 7.46 (1H, d,6.8), 6.86 (1H, d, *J*=6.8), 7.68 (2H, d, *J*=8.5), 7.68 (2H, d, *J*=8.5), 6.95(1H, d, *J*=10), 5.87 (1H, d, *J*= 12.5)

^13^C NMR (500 MHz, CDCl_3_): δ_C_ 175.6, 99.1, 103.0, 100.5, 115.9, 154.8, 115.9, 110.0, 64.2

#### Mixture of 7,8 -(3”,3”-dimethyl-pyrano)-4’-hydroxy flavonol and 4’,7,8-trihydroxy flavonol (3)

White powder, ^1^H NMR (500 MHz, CDCl_3_): δ_H_ 6.81 (1H, s, H-3), 7.46 (1H, d, *J*=7.2, H-5), 7.30 (1H, d, *J*=7.2, H-6), 8.00 (2H, d, *J*=6.8, H-2’), 6.88 (2H, d, *J*=6.8, H-3’), 6.88 (2H, d, *J*=6.8, H-5’), 8.00 (2H, d, *J*=6.8, H-6’)

^13^C NMR (500 MHz, CDCl_3_): δ_C_ 151.6, 140.3, 123.4, 152.9, 136.0, 151.6, 124.8, 127.2, 161.4, 127.2

#### Mixture of α-amyrin and β-amyrin (4)

**α-amyrin:** White powder, ^1^H NMR (500 MHz, CDCl_3_): δ_H_ 5.28 (t, 1H, H-12), 3.24 (dd, 2H, *J*=4.4, 4.8, H-3), 1.16 (s, 3H, H-27), 1.11 (s, 6H, H-26), 1.01 (s, 6H, H-28), 0.98 (s, 6H, H-25), 0.93 (d, 3H, *J*=3.2, H-30), 0.82 (d, 3H, *J*=6.5, H-29), 0.80 (s, 6H, H-23), 0.79 (s, 6H, H-24)

#### β-amyrin

White powder, ^1^H NMR (500 MHz, CDCl_3_): δ_H_5.31 (t, 1H, H-12), 3.24 (dd, 2H, *J*=4.4, 4.8, H-3), 1.28 (s, 3H, H-27), 1.11 (s, 6H, H-26), 1.01 (s, 6H, H-28), 0.98 (s, 6H, H-25), 0.95 (s, 6H, H-29,30) 0.80 (s, 6H, H-23), 0.79 (s, 6H, H-24)

#### Monoglyceride of Stearic Acid (6)

Colorless crystals, ^1^H NMR (500 MHz, CDCl_3_): **Glycerol backbone:** -CH_2_-O-CO: 4.195 (2H, dd, *J* = 13.6, 5.6 Hz), -CH: 3.957 (1H, m), -CH_2_: 3.732 (2H, dd, J = 11.6, 4.0 Hz), 3.617 (1H, dd, J = 11.6, 5.6 Hz). **Stearic acid part:** -CH2-CO-O: 2.37 (t, 2H), -CH2: 1.64 (d, 2H), 4-CH2: 1.31 (d, 8H), 10-CH2: 1.25 (s, 20H), -CH3: 0.90 (t, 3H)

**Figure 2:**
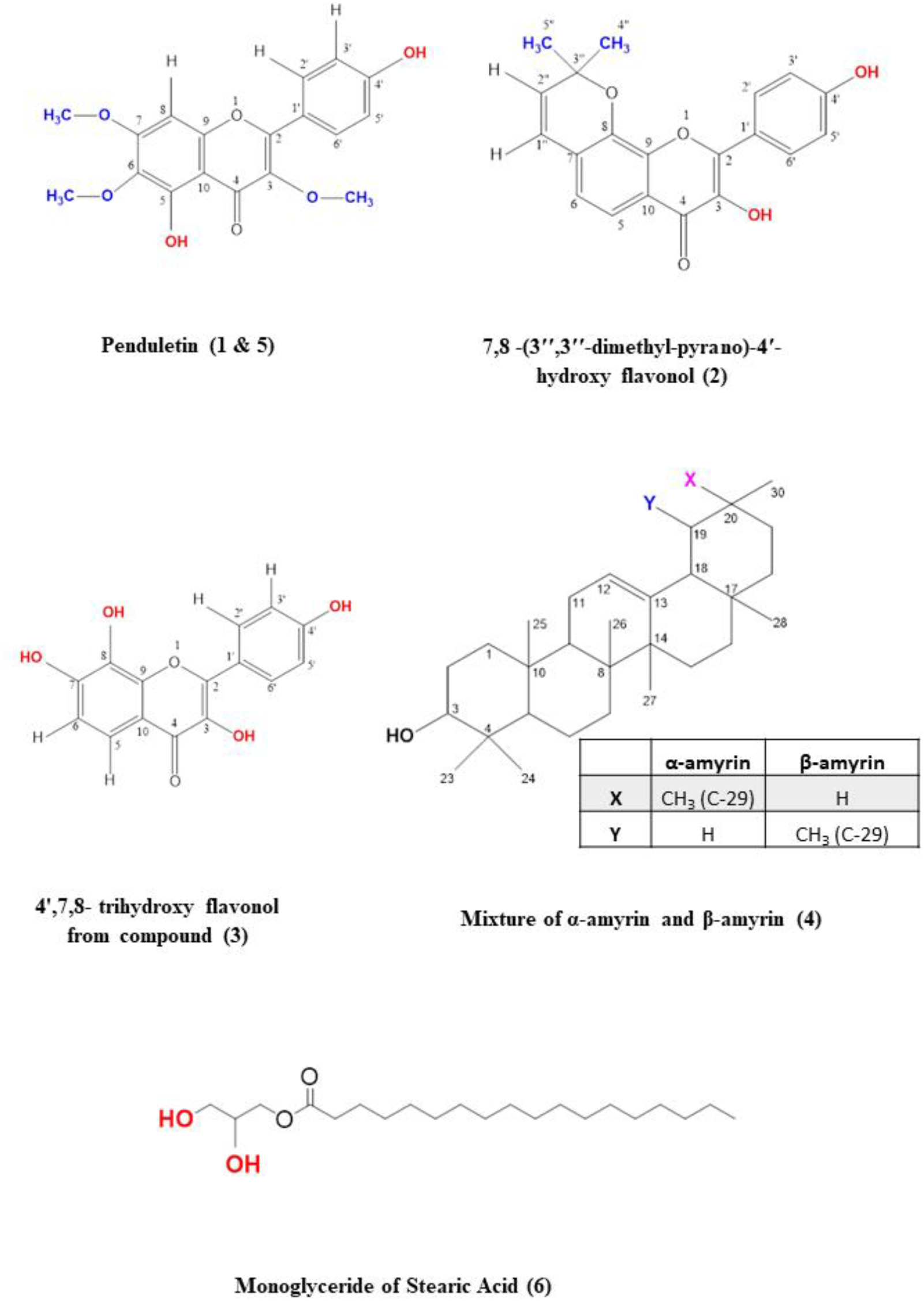
Isolated compounds from *Colocasia gigantea* and *Colocasia affinis*.

## Results and discussion

Dichloromethane fraction of both plants were used in phytochemical screening and compound isolation. A total of 3 compounds were elucidated from the dichloromethane soluble fraction of *C. affinis* as a result of consecutive chromatographic separation and purification. The compounds isolated from *C. affinis* are Penduletin (1), 7,8 -(3’’,3’’-dimethyl-pyrano)-4’-hydroxy flavonol (2) and mixture of 7,8 -(3’’,3’’-dimethyl-pyrano)-4’-hydroxy flavonol and 4’,7,8-trihydroxy flavonol (3). Another 4 compounds were isolated from the dichloromethane soluble fraction of *C. gigantea* following the same technique and those are mixture of α-amyrin and β-amyrin (3), penduletin (5) (which was isolated from *C. affinis* also) and monoglyceride of stearic acid (6).

The ^1^HNMR spectrum of compound 1 and 5 (Figure 3) in CDCL_3_ indicated the presence of 3-methoxy groups at δ_H_ 4.04 (3H, s), δ_H_ 4.02 (3H, s) and δ_H_3.90 (3H, s). The two paired of ortho coupled (*J= 7.2 Hz*) doublets as δ_H_7.04 and δ_H_ 7.90 showed that ring B is monosubstituted at C-4’. The peak at δ_H_ 12.79belongs to 5-OH. There is a one proton singlet at δ_H_ 6.59.There are 3 possible positions for this singlet which are C-3, C-6 and C-8. Another one proton singlet at δ_H_ 6.37 could be assigned to the -OH group at C-4’. All these ^1^H NMR data are in close agreement with the published value of Penduletin (17,18) where the singlet at δ_H_ 6.59 is at C-8 position and three methoxy groups are at C-3, C-6 and C-7 positions. Thus, the structure of compound 1 and 5 can be concluded as Penduletin.

**Figure 3:**
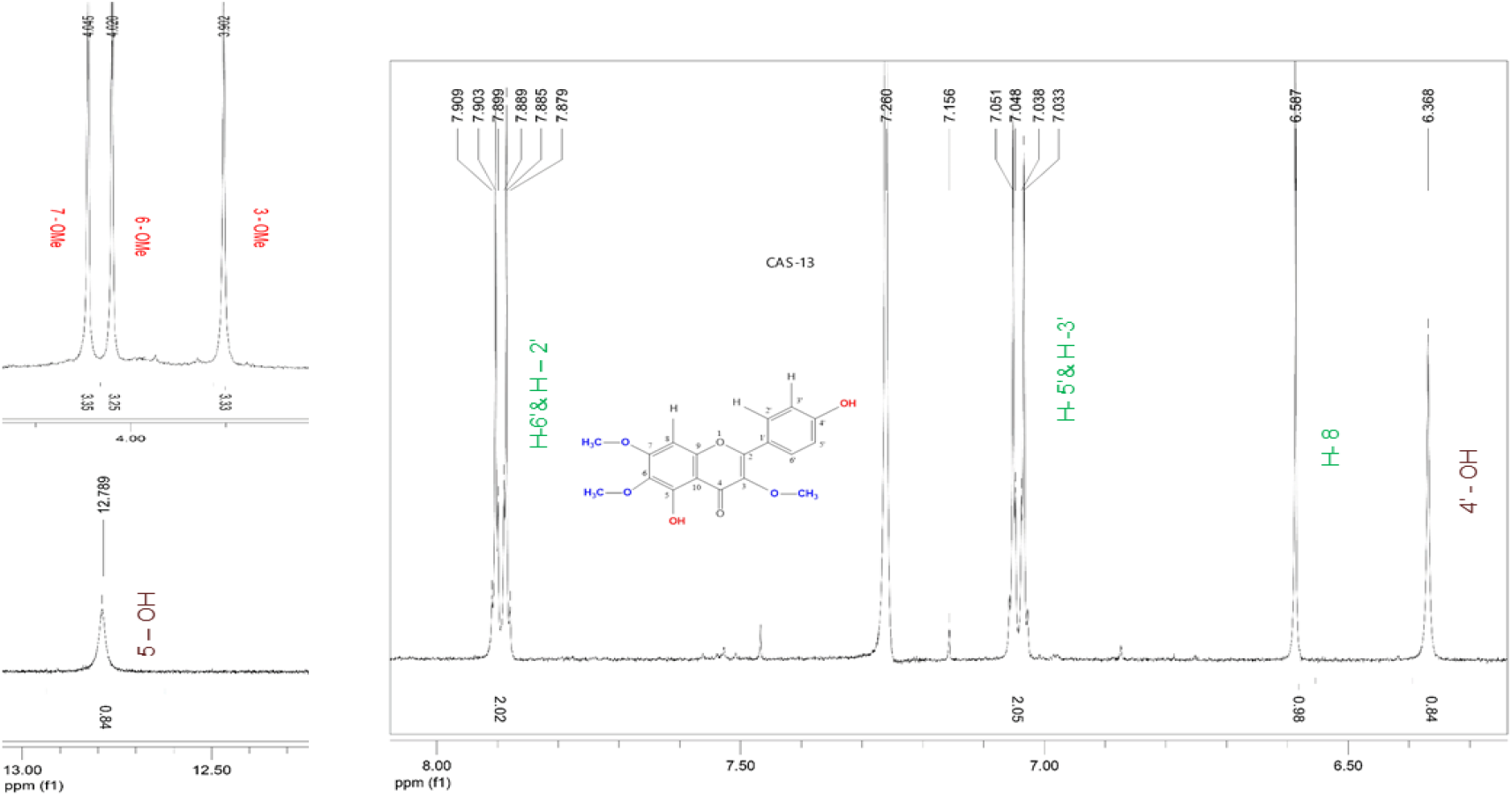
^1^H NMR spectrum of compound 1 and 5 as Penduletin.

The ^1^H NMR spectrum of compound 2 (Figure 4) showed two aromatic systems including one 1,4-disubstitued benzene ring with the doublet at δ_H_ 7.68 (2H, d,*J= 6.8 Hz*), one tetra-substituted benzene ring with the ^1^H signals at δ_H_ 6.86 (1H, d,*J= 6.8 Hz*), δ_H_7.46 (1H, d,*J= 6.8 Hz*) and a 3”,3”-dimethyl-pyrano group in AS-17. The presence of 3”,3”-dimethyl-pyrano group can be proved by two tertiary-methyl signals at δ_H_ 1.55 (6H, s) and two olefinic proton signals at δ_H_ 5.87 (1H, d,*J* =10 Hz) and δ_H_ 6.95 (1H, d, *J=10 Hz*). All these values of chemical shifts ae in close agreement with the published values of Citrusinol except the presence of an extra aromatic proton signal at which could be assigned to H-5. Another change in chemical shift of H-6 proton at δ_H_ 6.86 differs largely in values which is at δ_H_ 6.21 for Citrusinol (19). This high value of chemical shift can be justified only if the position of oxygen atom of pyran ring is at C-8 rather than C-7. Thus, the structure of compound 2 can be proposed as 7, 8-(3”,3”-dimethyl-pyrano)-4’-hydroxy flavonol.

**Figure 4:**
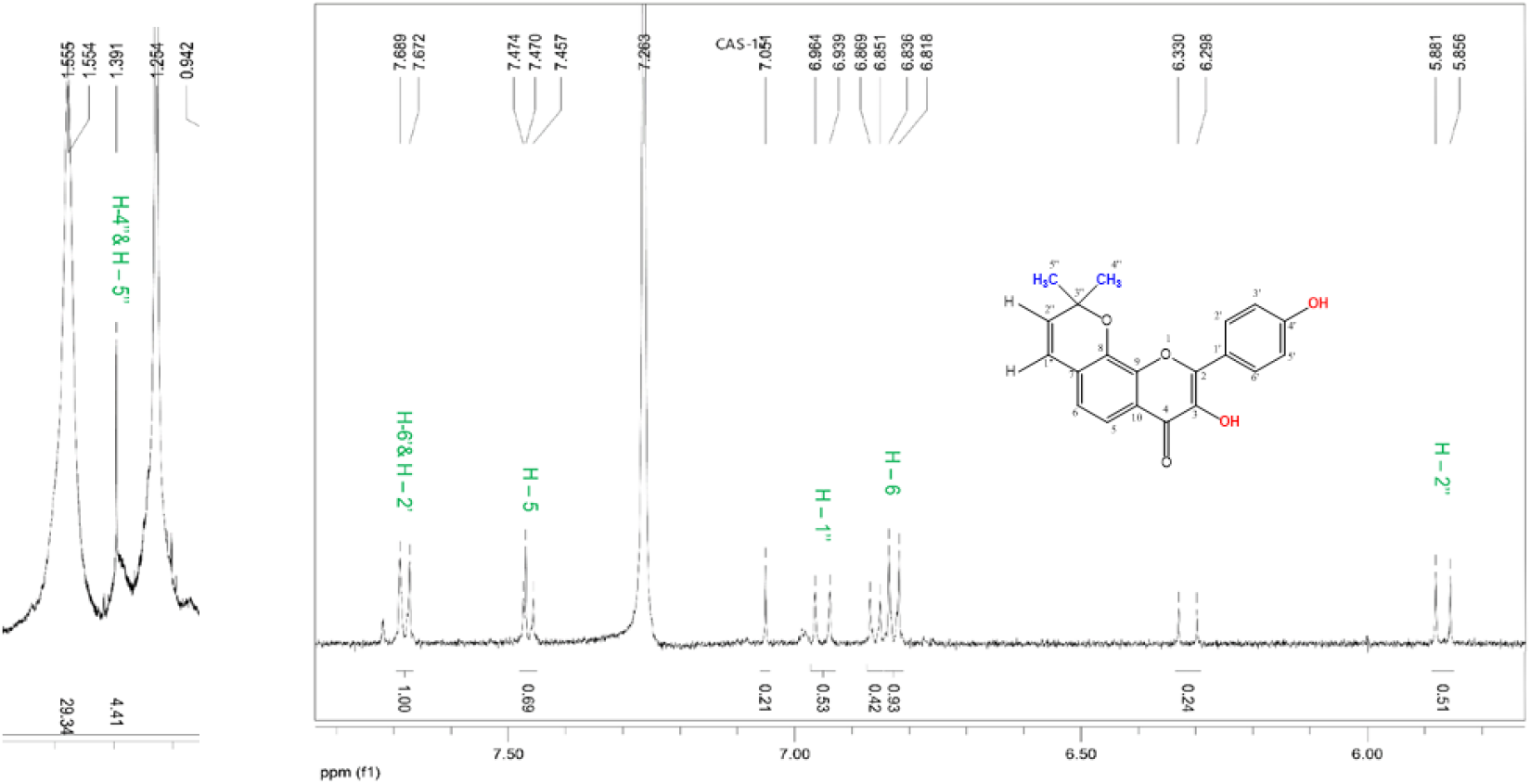
^1^H NMR spectrum of compound 2 as 8-(3”,3”-dimethyl-pyrano)-4’-hydroxy flavonol.

The ^1^H NMR spectrum (CDCl_3_, 400 MHz) of compound 3 show signals that exactly matches with that of compound 2 (7,8-(3’’,3’’-dimethyl-pyrano)-4’-hydroxy flavonol). But the presence of additional peaks suggests that the presence of another compound as mixture. The additional signals include two para-coupled doublets with *J=6.8Hz* at δ_H_8.00 and δ_H_.88, each integrating for two protons which were assigned to the coupled H-2’ and H-6’, and H-3’ and H-5’ respectively. A typical ortho-coupled signal for protons at C-5 and C-6 at δ_H_7.46 (1H,d,*J= 7.2Hz*’ and at δ_H_7.30 (1H,d,*J= 7.2Hz*).All these additional proton signal values are in close agreement with that of published values (20) with the structure (Figure 5) and characterized as 4’,7,8-trihydroxy flavonol. ^13^C NMR also supports the argument. Thus, the structure of compound 3 appeared as a mixture of 4’,7,8-trihydroxy flavonol with 7,8-(3”,3”-dimethyl-pyrano)-4’-hydroxy flavonol (compound 2) in a ratio of 1:2 according to peak height in ^1^H NMR spectrum.

**Figure 5:**
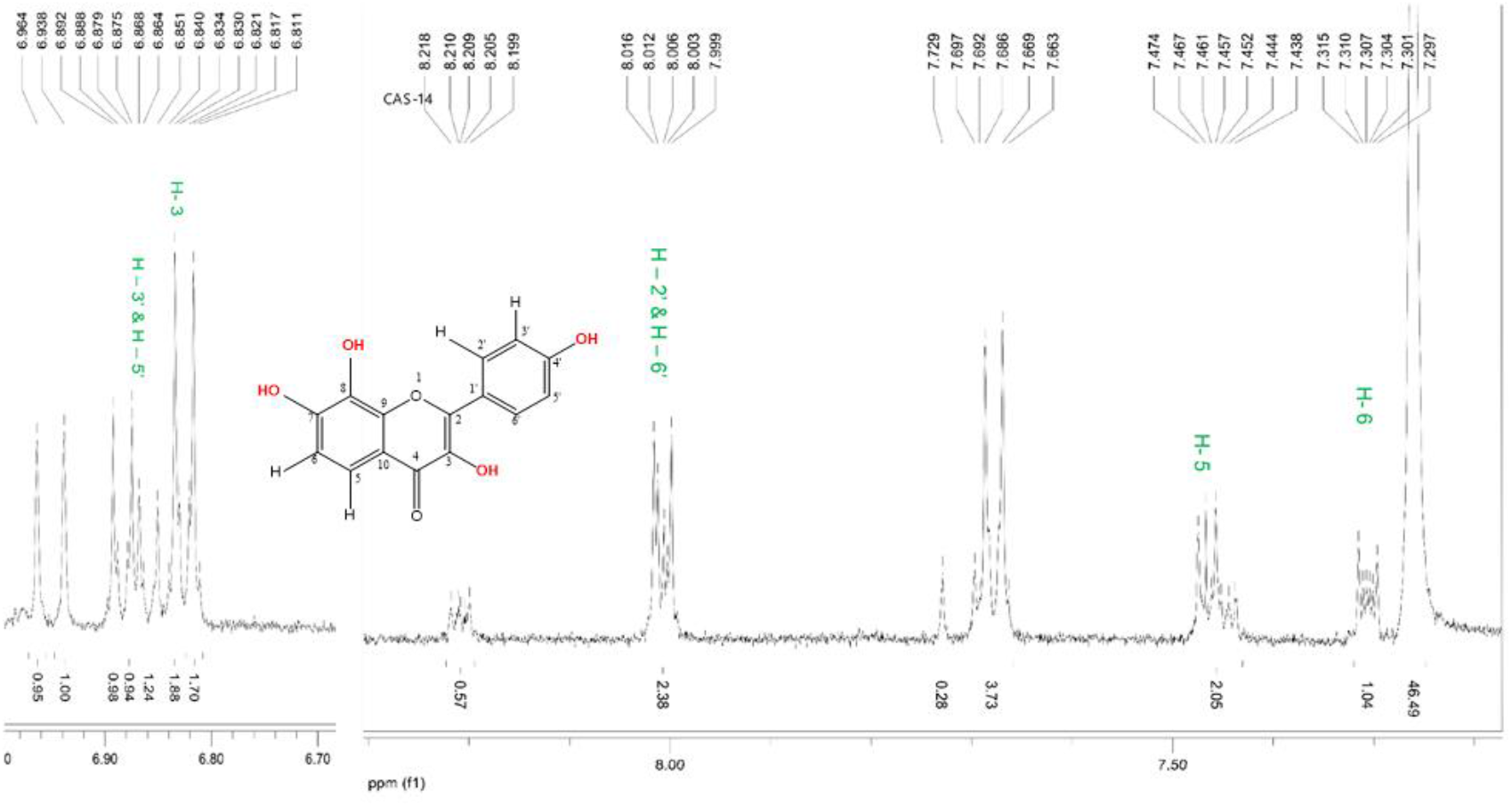
^1^H NMR spectrum of 4’,7,8-trihydroxy flavonol from compound 3.

The ^1^H NMR spectrum (400 MHz, CDCl_3_) of compound 4 (Figure 6) showed two triplets at δ5.32 ppm and 55.28 ppm characteristic of olefinic proton (H-12) of α- and β-amyrin respectively. Apart from that, eight singlet and two doublets of nCH3 protons were identified in the range δ_H_1.28-0.79 ppm. The singlet at δ_H_0.89 (3H) and δ_H_ 0.88 ppm (3H) indicated the presence of CH3-28. That means that the analyzed component was not an acid with the carboxylic group at C-17. The peaks of the methyl groups of α-amyrin were identified at δ_H_1.16 ppm (singlet; CH_3_-27), δ_H_0.93 ppm (doublet-doublet; *J= 3.2 Hz*; CH_3_-30) and δ_H_0.82 pm (singlet; CH_3_-29). The protons of CH_3_-27, CH_3_-29 and CH_3_-30 groups of β-amyrin had peaks at δ_H_1.28 ppm (singlet; 3H; CH_3_-27), δ_H_0.95 ppm (singlet; 6H; CH_3_-29 and 30). The other signals were identical for both amyrins: δ_H_1.11 (6H, s; CH_3_-26), δ_H_1.01 (6H, s; CH_3_-28), δ_H_0.98 (6H, s; CH_3_-25), δ_H_0.82 (6H, s; CH_3_-23), δ_H_0.80 (6H, s; CH_3_-24). The above data enabled identification of compound 4 as a mixture of α- and β-amyrin with the ratio of 4:3 based on the ^1^H NMR peaks. This identity was further confirmed by direct comparison of its ^1^H NMR spectrum with that recorded for ^1^H NMR of mixer of α-amyrin and *β-* amyrin in CDCl_3_ (500 MHz, CDCl_3_) (21).

**Figure 6:**
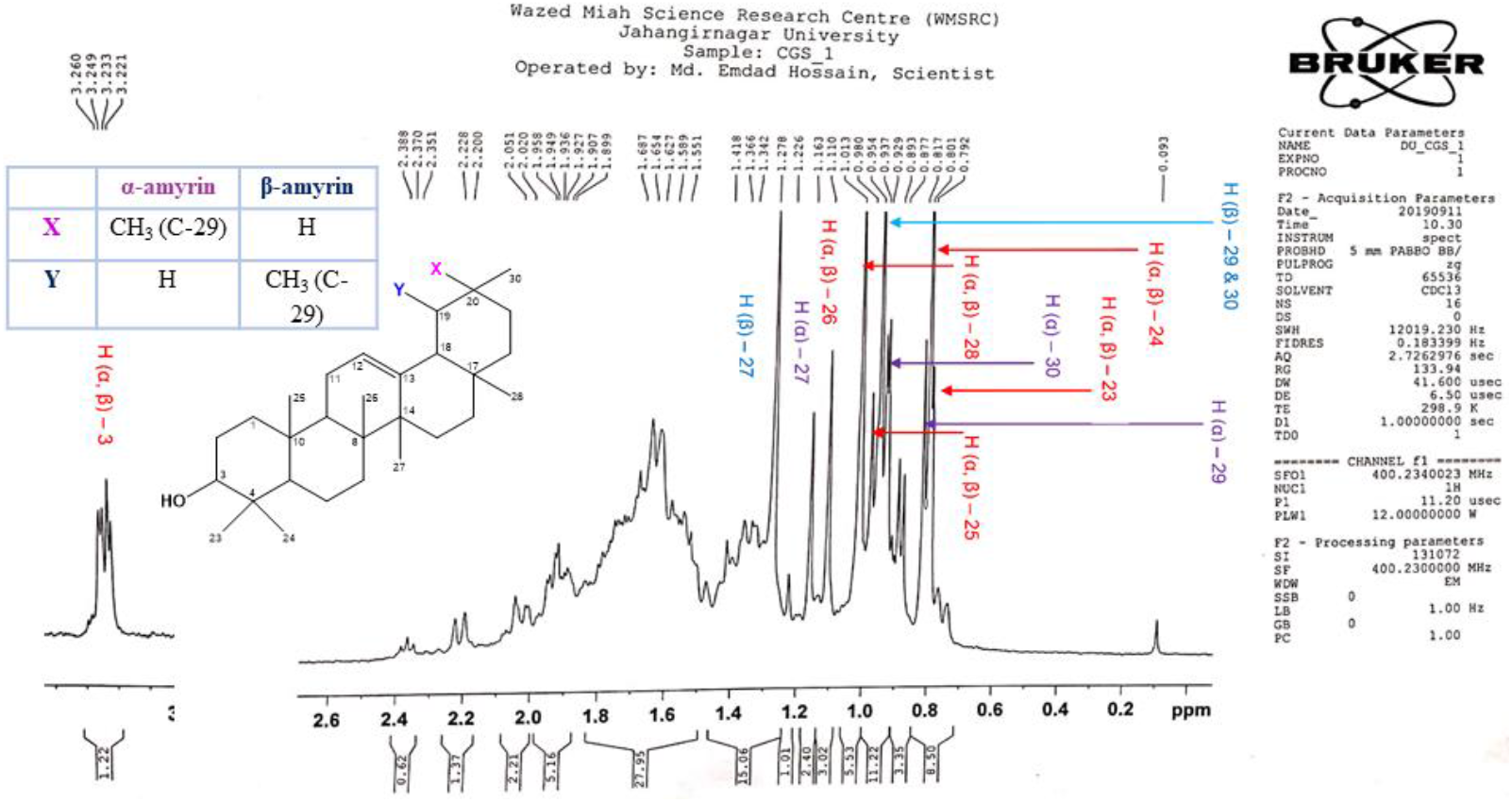
^1^H NMR spectrum of compound 4 as mixture of α-amyrin and β-amyrin.

The structure of compound 6 can be proposed as the monoglyceride of saturated fatty acid. The ^1^H NMR (400 MHz, CDCl_3_) spectrum showed the signals at δ_H_ 4.195 (2H, dd, *J* = 13.6, 5.6 Hz), δ_H_ 3.957 (1H, m) and δ_H_ 3.732 (2H, dd, *J* = *11.6, 4.0* Hz). This group of signals can be attributed for the protons of the (-CH_2_-CH-CH_2_-) backbone (glycerol structure). The upfield peak at 5 0.903 with three proton intensity corresponds to one −CH_3_ protons. Two additional distinct group of upfield signals at δ_H_ 2.374 (2H, t, *J* = 7.6 Hz), δ_H_ 1.64 (2H, m), δ_H_ 1.313 (8H, m) and δ_H_ 1.25 (20H, m) could correspond to 16 (sixteen) methylene groups. The presence of an additional oxymethine proton in the carbon chain can be justified by the signal at δ_H_ 3.617 (1H, dd, *J* = 11.6, 5.6 Hz). Comparing all the spectral data the structure of the compound 5 with previously documented study (22) can be concluded as monoglycerides of stearic acid with the structure drawn in Figure 7.

**Figure 7:**
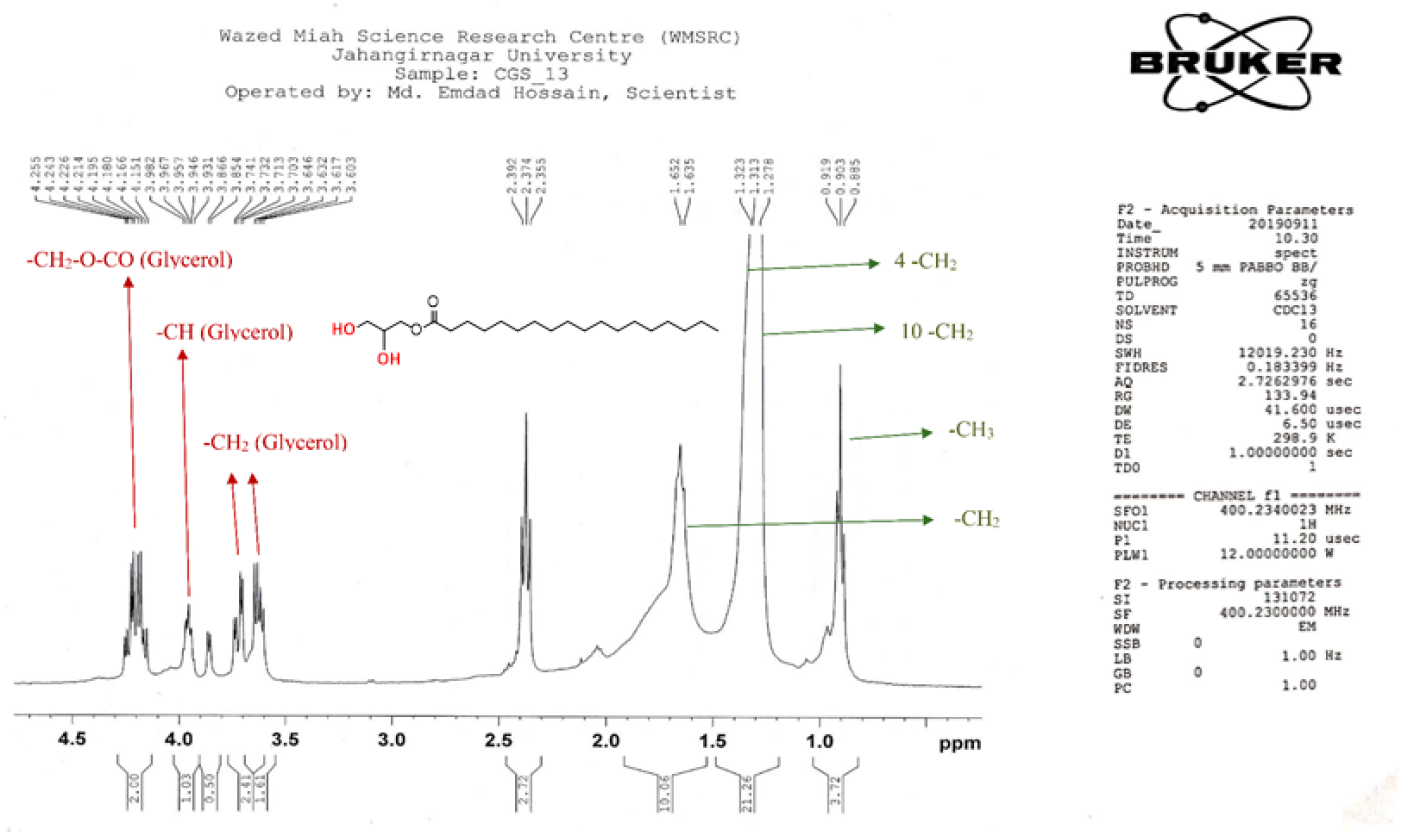
^1^H NMR spectrum of compound 6 as Monoglyceride of Stearic Acid.

## Conclusion

Seven compounds were isolated from two plants among which penduletin was found in both species. On the basis of these spectral data, the structures of the isolated compounds were characterized and it is visible that both plants contain several types of flavonoids and due to this these *Colocasia* plants can be considered as well spring of bioactive phytoconstituents to establish new drug molecules and deserve more intensive researches.

## Acknowledgements

The authors are grateful to Department of Pharmaceutical Chemistry, University of Dhaka

## Authors Contribution

All authors contributed equally.

## Availability of data and materials

Not applicable.

